# Muscle contractile properties directly influence shared synaptic inputs to spinal motor neurons

**DOI:** 10.1101/2023.11.30.569389

**Authors:** Hélio V. Cabral, J Greig Inglis, Alessandro Cudicio, Marta Cogliati, Claudio Orizio, Utku Yavuz, Francesco Negro

**Author notes:** Corresponding author: Prof. Francesco Negro, Department of Clinical and Experimental Sciences, Università degli Studi di Brescia, Viale Europa 11, Brescia, 25121, Italy.

## Abstract

Alpha band oscillations in shared synaptic inputs to the alpha motor neuron pool can be considered an involuntary source of noise that hinders precise voluntary force production. This study investigated the impact of altering muscle length on the shared synaptic oscillations to spinal motor neurons, particularly in the physiological tremor band. Fourteen healthy individuals performed low-level dorsiflexion contractions at ankle joint angles of 90° and 130°, while high-density surface electromyography (HD-sEMG) was recorded from the tibialis anterior (TA). We decomposed the HDsEMG into motor units spike trains and calculated the motor units’ coherence within the delta (1-5 Hz), alpha (5-15 Hz) and beta (15-35 Hz) bands. Additionally, torque steadiness and torque spectral power within the tremor band was quantified. Results showed no significant differences in torque steadiness between 90° and 130°. In contrast, alpha band oscillations in both synaptic inputs and force output decreased as the length of the TA was moved from shorter (90°) to longer (130°), with no changes in delta and beta bands. In a second set of experiments, evoked twitches were recorded with the ankle joint at 70° and 130°, revealing longer twitch durations in the longer muscle lengthen condition compared to the shorter. These experimental results, supported by a simple computational simulation, suggest that increasing muscle length enhances the muscle’s low-pass filtering properties, influencing the oscillations generated by the Ia afferent feedback loop. Therefore, this study provides valuable insights into the interplay between muscle biomechanics and neural oscillations.

## Introduction

The generation of voluntary movement relies on a sequence of integrated processes that culminate in the production and modulation of a muscle force output (Heckman and Enoka, 2012, Enoka and Farina, 2021). The pool of alpha motor neurons plays a pivotal role in these complex processes by integrating common and independent synaptic inputs into individual motor unit action potential train outputs which are subsequently propagated to the active muscle (Ishizuka et al., 1979, Lemon, 2008, Duchateau and Enoka, 2011). Rather than acting independently, the resultant discharge times of individual motor units exhibit similar behaviours, as evidenced by the high correlation between their activities (Sears and Stagg, 1976, Kirkwood and Sears, 1978, De Luca et al., 1982, Datta and Stephens, 1990, Farmer et al., 1993, Hug et al., 2023). These common fluctuations observed in the output of neuronal ensembles have been attributed to shared synaptic projections across motor neuron pools, which may have cortical or spinal origins (Nordstrom et al., 1992, Farmer et al., 1993, Bremner et al., 1991, Negro and Farina, 2011, Farina et al., 2014). Given the main role of cortical pathways in the voluntary control of movement, shared projections from the motor cortex are believed to be the primary source of correlation between motor neuron spike trains, particularly in the delta (associated with voluntary force corrections) and beta bands (linked with corticomuscular coherence) (Kirkwood and Sears, 1978, Datta and Stephens, 1990, Datta et al., 1991, Baker et al., 1997, Conway et al., 1995, Bräcklein et al., 2022). These shared projections hold an important functional role, as the synergistic activation of multiple motor neurons by these common oscillations facilitates the optimization of the motor control system (d’Avella and Bizzi, 2005, Berniker et al., 2009, Laine et al., 2015).

The shared cortical projections are not the exclusive source of correlations between motor unit spike trains in human muscles. Notably, during isometric voluntary contractions, the force output exerted includes a significant portion of involuntary rhythmic fluctuations in the frequency range of 5-15 Hz (alpha band), often associated with physiological tremor (Lippold, 1971, McAuley and Marsden, 2000). These oscillations, resulting from physiological tremor, can be partially attributed to afferent inputs originating from the active muscles themselves, which project to the motor neuron pool (Lippold, 1970, Hagbarth and Young, 1979, Cresswell and Löscher, 2000, Christakos et al., 2006, Laine et al., 2016). More specifically, the spinal stretch reflex establishes a feedback loop that functions as a servomechanism system aimed at maintaining stability at the muscle level (Lippold, 1971). Due to inherent delays in signal transmission within this feedback loop, oscillations at resonant frequencies, which were not present in the initial input of the motor neuron pool, tend to emerge (Halliday and Redfearn, 1956, Lippold, 1971, Lippold, 1970). Consequently, the resonance behaviour can lead to oscillations at these frequencies in the neural drive to the active muscle, resulting in variations in steadiness of the muscle force output (McAuley and Marsden, 2000, Inglis and Gabriel, 2021). In this context, physiological tremor can be seen as a source of common noise that interferes with precise voluntary force production. Previous studies have suggested that increased fluctuations in force output steadiness is likely the result of common noise in the neural commands sent to active motor units (Feeney et al., 2018, Negro et al., 2009, Thompson et al., 2018). Therefore, understanding the mechanisms involved in the modulation of alpha band oscillations (physiological tremor) is extremely important in the investigation of neuromuscular control.

Recent investigations have demonstrated that oscillations in force output within the alpha band can be modulated by a range of factors. These factors include changes in nociceptive input (Yavuz et al., 2015), enhancement of serotonin availability by pharmacological means (Henderson et al., 2022), and alterations in the demands of a visuomotor task (Laine et al., 2014). Of particular interest are changes in muscle length which have been shown to exert a direct influence on alpha oscillations in force output, with shorter muscle lengths being associated with an increase in physiological tremor (Jalaleddini et al., 2017). A potential explanation for this phenomenon, derived from computational simulations, is related to alterations in the modulation of the gamma static fusimotor drive following changes in muscle length (Jalaleddini et al., 2017). Additionally, these changes may occur as a result of mechanistic changes at the periphery, such as, the amount of musculoskeletal stiffness or joint laxity (Clamann and Schelhorn, 1988, Rack and Westbury, 1969, Powers and Binder, 1991). Furthermore, altering the muscle length affects the average twitch profile of motor units, resulting in changes to the muscle’s low-pass filtering properties when converting motor unit spike trains into mechanical movements about a joint (Rack and Westbury, 1969, Mannard and Stein, 1973, Bawa and Stein, 1976). Specifically, longer muscle lengths result in longer twitch durations, subsequently intensifying the low-pass filtering effect of the neural drive to the muscle. The enhanced effect of filtering the averaged motor unit twitch, in turn, may directly influence the behaviour of the afferent loop, leading to modulated oscillations in physiological tremor. However, whether this phenomenon occurs remains unexplored in the literature and, to date, no previous study has experimentally investigated the effect of altered muscle length on alpha oscillations in common synaptic inputs to the alpha motor neuron pool.

In the present study, we aimed to investigate whether alterations in muscle length could influence the common synaptic oscillations to spinal motor neurons, particularly in the tremor band (5-15 Hz). To achieve this, we decomposed motor unit spike trains from high-density surface electromyograms recorded from the tibialis anterior (TA) at two different muscle lengths. Additionally, to interpret the results derived from the motor unit spike trains, we conducted a second set of experiments, combined with computational simulations, to investigate how changes in muscle length affect the low-pass characteristics of the TA’s electrically evoked potentials (twitches).

## Methods

### Participants

Fourteen healthy individuals (1 female; mean ± SD: age 26 ± 3 years; height 178 ± 9 cm; mass 76 ± 10 kg) volunteered to participate in this study. All participants were free of any neuromuscular abnormalities and had no history of lower limb injury or lower leg pain that would impact their ability to produce voluntary contractions. All participants provided written informed consent before starting the experiments. This study was conducted in accordance with the latest version of the Declaration of Helsinki and approved by the local ethics committee (code NP2490).

### Experimental protocol

The study consisted of a single experimental session lasting ∼1h. Participants were comfortably seated with their right leg placed on a custom-built jig designed to isolate the ankle joint during dorsiflexion contractions. The right knee was fully extended, the hip was flexed at 70° (0° being the hip fully extended) and the right foot fixed with straps to an adjustable footplate that held the ankle at the specified joint angles. This footplate was connected to a load cell (SM-500 N, Interface, Arizona, USA) to record the dorsiflexion isometric torque produced by the dorsiflexors, more specifically the TA (**Figure 1A**). Additional straps were placed around the thigh and knee to ensure the torque produced relied solely on the dorsiflexor muscles. The subjects were asked to keep their left leg straight and relaxed on the side of the jig.

**Figure 1:**
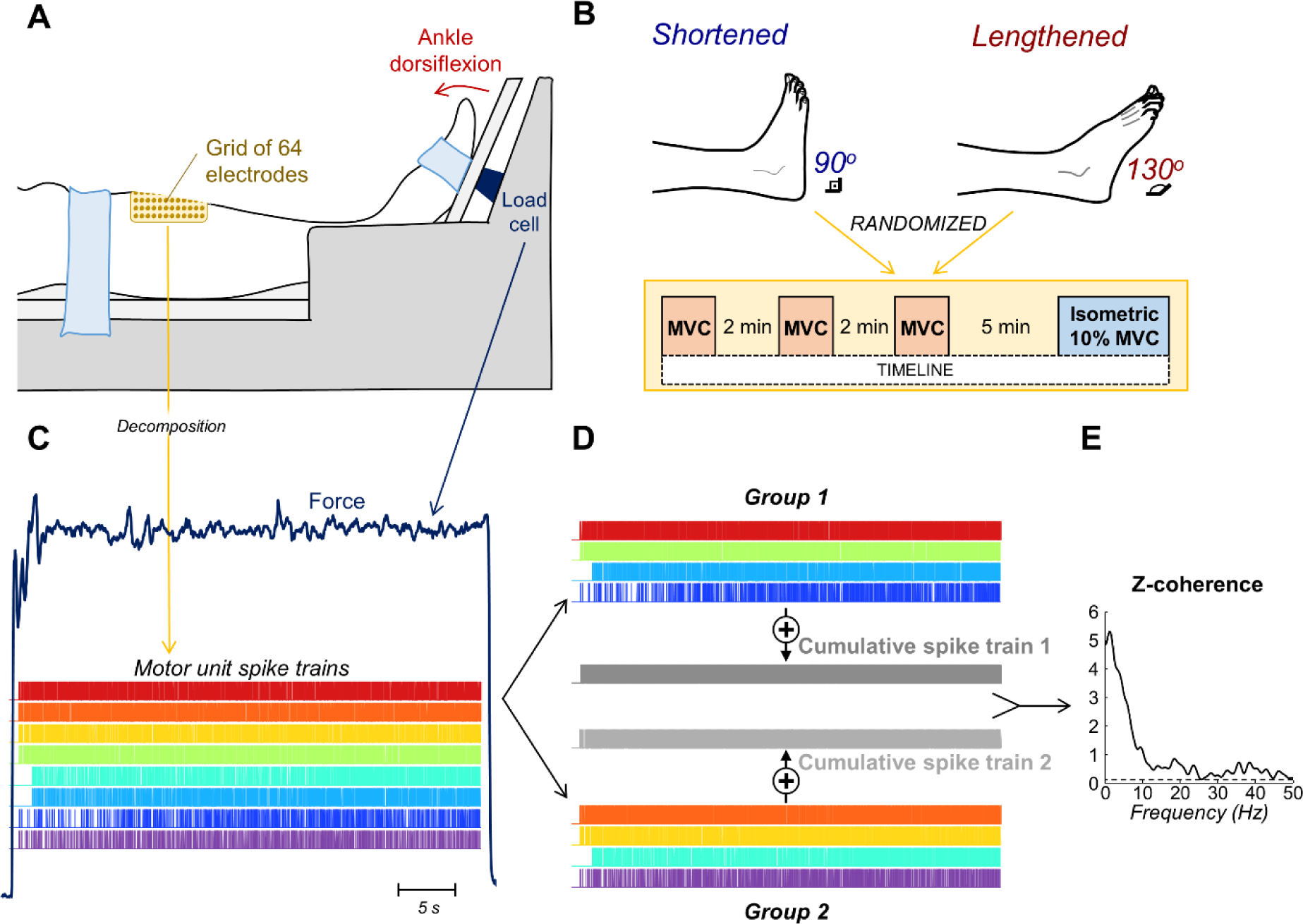
Experimental setup and motor unit analysis. A, Schematic representation of the participants’ foot position for measuring dorsiflexion isometric torque. High-density surface electromyograms (HDsEMG) were acquired from the tibialis anterior (TA) muscle using a grid of 64 electrodes. B, Ankle joint angles selected to represent conditions where the TA muscle was shortened (90°) and lengthened (130°). The tasks performed for each angle are shown in the yellow box. C-E, Steps involved in the motor unit analysis. After decomposition from the HDsEMG recordings (C), the cumulative spike trains of two equally sized groups of motor units were obtained (D), and the coherence between them was calculated (E).

#### Voluntary contractions

All experimental tasks were repeated for two different ankle joint angles: 90° (0° of plantar flexion) and 130° (40° of plantar flexion). These angles were selected to represent conditions in which the TA muscle was shortened (90°) and lengthened (130°) (top part of **Figure 1B**). Condition order was randomized across participants and a minimum of ten min of rest was provided between conditions. For each condition, participants repeated the same experimental protocol (bottom part of **Figure 1B**). Each participant performed three voluntary maximum isometric contractions (MVCs) for three seconds (s), with a two-minute (min) rest between each bout. The greatest value across the three MVCs was considered as the maximal isometric dorsiflexion torque and used as a reference to set the target of the submaximal force outputs of the corresponding condition. Following a short familiarization period where the participants practiced the task to ensure they were able to match the level of submaximal torque output, participants were instructed to perform an isometric dorsiflexion contraction following a trapezoidal profile. For both conditions (90° and 130°), the profile involved a linear increase at a rate of 5% MVC/s, a plateau at 10% MVC for 40 s, and a linear decrease at a rate of 5% MVC/s. Visual feedback from the target and produced dorsiflexion torque output was displayed on a computer monitor positioned ∼1m in front of the participant.

#### Evoked contractions

To assess the effect of muscle length changes on the TA twitch characteristics, we conducted a second set of experiments in a subgroup of five participants. On a separate day from the initial experiments, participants returned to the laboratory, and electrical stimulation was applied to the common peroneal nerve of their right leg while they were positioned as described in the first set of experiments. First, a round cathode electrode (20 mm diameter; Spes Medica, Genova, Italy) filled with conductive paste was placed on the skin posterior and inferior to the head of the fibula and a rectangular anode electrode (size 50 x 100 mm; UltraStim, Axelgaard Manufacturing, CA, USA) was positioned on the opposite side of the knee. The posterior inferior region to the head of the fibula was identified through palpation by an experienced researcher. Once the stimulating electrode was fixed in position, the current intensity leading to the largest evoked dorsiflexion torque output response was defined using a staircase current profile. Specifically, a train of rectangular pulses (200 µs of duration; 1 Hz frequency) with increasing amplitudes (staircase protocol) was delivered until no clear increase in dorsiflexion torque output could be visualized; this level was defined as the maximal stimulation intensity. After a 90 s period of rest, rectangular pulses (200 µs of duration; 1 Hz frequency) were applied to electrically evoke twitches from the TA for two distinct ankle joint positions: 70° (20° of dorsiflexion) and 130° (40° of plantar flexion). These two extreme ankle joint angles were chosen to further investigate the effect of biomechanical changes about the ankle joint and muscle length on the filter characteristics of muscle twitch force. When the ankle joint was placed at an angle of 70° this shortened the TA muscle length, while at 130° the TA muscle length was lengthened. For each condition, fifteen supramaximal evoked potentials (20% over the maximal stimulation intensity defined during the staircase protocol) where delivered to the participant. The order of ankle joint angle was randomized with 90-s rest between each interval.

### Data collection

High-density surface electromyograms (HDsEMG) were acquired from the TA during the submaximal isometric task using a two-dimensional adhesive grid of 64 electrodes (8 mm inter-electrode distance; OT Bioelettronica, Turin, Italy). The grid was positioned longitudinally on the TA muscle belly, which was determined via palpation (**Figures 1A**). Prior to electrodes placement, the area was shaved, mildly abraded (EVERI, Spes Medica, Genova, Italy) and cleaned. The electrode-skin contact was ensured by filling the foam cavities with conductive paste (AC cream, Spes Medica, Genova, Italy). The reference electrode was positioned on the right ankle at the level of the malleoli. HDsEMG and torque outputs were digitized synchronously at a sampling frequency of 2048 Hz using a 12-bit A/D converter (10-500 Hz bandwidth; EMG-USB2+, OT Bioelettronica, Turin, Italy). HDsEMG signals were recorded in monopolar derivation and amplified to maximize signal resolution while avoiding saturation.

### Data analysis

Torque output, HDsEMG and evoked twitches were analysed offline using MATLAB (version 2022b) custom-written scripts.

### Torque Output

Torque outputs acquired during submaximal dorsiflexion contractions were low-pass filtered at 15 Hz using a third-order Butterworth filter. The coefficient of variation (CoV) was estimated to quantify the variation of force output at each ankle joint angle (90° and 130°). To this end, a stationary portion of the force signal was chosen from the 40-s plateau region. The CoV was computed for 30-s moving windows with one sample step. Additionally, the power spectral density of torque output was estimated using Welch’s method (*pwelch* function in MATLAB; 1-s Hanning windows with 1948 samples of overlap). Then, for each condition, the mean power within the alpha band (5-15 Hz) was calculated and retained for further analysis. For this analysis, the central 30-s of the torque output was used for all participants and conditions.

### HDsEMG decomposition and estimates of common synaptic input

The steps involved in the HDsEMG analysis are schematically represented in **Figures 1C-E**. First, monopolar HDsEMG signals were bandpass filtered with a third-order Butterworth filter (20-500 Hz cut-off frequencies). After visual inspection, channels with low signal-to-noise ratio or artifacts were discarded. The HDsEMG signals were then decomposed into their constituent motor unit spike trains using a convolutive blind-source separation algorithm (Negro et al., 2016). Briefly, after extending and whitening HDsEMG signals, a fixed-point algorithm that seeks sources that maximize a measure of sparsity was applied to identify the sources (i.e., motor unit spike trains; **Figure 1C**). The spikes were separated from the noise using K-means and, while iteratively updating the motor unit separation vectors, the discharge times estimation was further refined by minimizing the coefficient of variation of the inter-spike intervals. This decomposition method has been extensively applied to assess the activity of single TA motor units (Negro et al., 2016, Cogliati et al., 2020, Cudicio et al., 2022). After the automatic identification of motor units, missed or misidentified motor unit discharges which produced non-physiological discharge rates were manually identified and iteratively edited by an experienced operator, and the subsequent motor unit spike trains were re-estimated (Martinez-Valdes et al., 2017, Hassan et al., 2020). This approach has been shown to be highly reliable across operators (Hug et al., 2021).

All the motor units decomposed from the experimental data were examined during the 30-s window centred midway through the torque output plateau. This section was chosen to exclude periods of torque output transition at the beginning and end of the trapezoidal contraction (McManus et al., 2019). In addition, motor units were discarded from the analysis when they did not discharge continuously for at least 20-s in the selected region. Coherence analysis was used to estimate the level of common synaptic input to the motor neuron pool (Negro and Farina, 2012, Castronovo et al., 2015a, Rossato et al., 2022). Coherence was estimated between cumulative spike trains (CST), each comprised of a minimum of two motor unit spike trains. Therefore, only participants with at least four motor units identified in each condition were included in the coherence analysis. First, the identified motor units from each condition were randomly divided into two equally sized groups (**Figure 1D**). The number of motor units included in each group was half of the minimal number of motor units identified across conditions. Thus if 18 and 12 motor units were detected for the 90° and 130° angles respectively, 6 motor units were included in each group. The discharge times in each group were then summed to obtain two CSTs (**Figure 1D**). Coherence was calculated between the two detrended CSTs using Welch’s periodogram with a 1-s Hanning window with an overlap of 95% (**Figure 1E**). This procedure was repeated for up to 100 randomly chosen combinations of two groups of motor units, and the average of all permutations was calculated (i.e., pooled coherence) (Castronovo et al., 2015b, Rossato et al., 2022). The Fisher z-transform was then applied to the pooled coherence estimates using an approximation for coherence calculations with overlapping windows (Gallet and Julien, 2011). Only z-scores greater than the confidence level were considered for further analysis. The confidence level was defined for each participant as the mean value of z-scores between 250 and 500 Hz, as no significant coherence is expected in this frequency range (Rossato et al., 2022). To investigate changes in common synaptic input between different TA muscle lengths, the averages of z-coherence within delta (1-5 Hz), alpha (5-15 Hz) and beta (15-35 Hz) bands were calculated. For purposes of visualization only (no statistical analysis), the z-coherence values were combined across all participants according to Baker et al. (2003).

### Tibialis anterior evoked twitches

Torque outputs acquired during electrically evoked contractions were triggered and averaged at the stimulation times to obtain an averaged twitch torque for each ankle joint angle (70° and 130°). The following variables were then extracted from the averaged twitch torque profiles: time to peak (*t_peak_*), measured as the time elapsed from the first detectable increase in torque to the peak twitch; and time to half-peak (*t_50%peak_*), determined as the time from the first detectable increase in torque to 50% of decay from the peak twitch. Considering the muscle acts as a low-pass filter when converting the motor unit spike trains into mechanical movement about a joint (Mannard and Stein, 1973, Bawa and Stein, 1976), to explore how the low-pass filter characteristics of TA changed during the different muscle lengths, we computed the normalized magnitude of the frequency-response of the twitch torque output obtained for each ankle joint angle (70° and 130°). The magnitude of the frequency-response represents the gain of the system as a function of frequency. We then calculated the cut-off frequency (*f_co_*) in which the gain was attenuated by -3dB.

### Simulations

To further investigate how the contractile properties of motor units influence the transmission of alpha band oscillations into force output, we used a well-established model that simulates the sequence of events from the excitation of an ensemble of motor neurons to the generation of isometric force output. This model, initially proposed by Fuglevand et al. (1993), has since been utilized in various studies (Taylor et al., 2002, Dideriksen et al., 2012, Dideriksen and Negro, 2018, Dideriksen et al., 2022, Gogeascoechea et al., 2023). The motor neuron parameters were selected based on an exponential distribution across the pool of motor neurons (Fuglevand et al., 1993). Similar to the number of motor units in the TA (Feinstein et al., 1955), the number of motor units in the model was set to 450, of which only those with a minimum discharge rate of 8 pulses per second were fully recruited (Taylor et al., 2002). The input to the motor neuron pool was modelled as a linear summation of common synaptic input to all motor neurons (low-pass filtered Gaussian noise < 15 Hz) and an independent noise input specific to each motor neuron (low-pass filtered Gaussian noise < 50 Hz). The model was implemented in MATLAB (The MathWorks Inc., Natick, Massachusetts, USA), and simulations were conducted with a sampling frequency of 1,000 Hz.

To assess the effect of motor unit twitch duration in the transmission of common synaptic noise into force output, we created two distinct models, model 1 and model 2. Specifically, we simulated an increase in the twitch contraction times of individual motor units by modifying the longest contraction duration for the motor unit pool from 80 to 140 ms (parameter *T_L_* in equation 14 of Fuglevand et al. (1993)). This alteration resulted in a median increase of ∼35 ms in the contraction time of motor unit twitches between the two models. The model with greater contraction time values (model 2) was used to simulate the muscle lengthening condition. Each model was repeated 10 times, as has been done in previous simulation studies (Dideriksen and Negro, 2018, Dideriksen et al., 2022). For each time, we computed the average z-coherence of motor units within delta (1-5 Hz), alpha (5-15 Hz) and beta (15-35 Hz) bands, following the same approach used on the experimental data.

### Statistical analysis

All statistical analyses were performed in R (version 4.2.2), using the RStudio environment. To compare the peak MVC, torque output steadiness (i.e., coefficient of variation of the torque output) and mean torque output power between conditions Wilcoxon signed-rank tests were used.

Linear mixed-effect models (LMM) were applied for all statistical analyses of motor unit data, as they account for the non-independence of data points within each participant. For both experimental and simulated data, the z-coherence averages were compared, separately for delta, alpha and beta bands, using random intercept models with ankle joint angle (90° and 130°) as the fixed effect and participant as the random effect (i.e., *z-coherence average ∼ 1 + condition + (1 | participant)*). LMMs were implemented using the package *lmerTest* (Kuznetsova et al., 2017) with the Kenward-Roger method to approximate the degrees of freedom and estimate the *p*-values. The *emmeans* package was used, when necessary, for multiple comparisons and to determine estimated marginal means with 95% confidence intervals (Lenth et al., 2019). The variables calculated from TA twitch torque outputs (*t_peak_*, *t_50%peak_* and *f_co_* of magnitude response) were presented descriptively for each of the five participants. To compare the motor unit twitch contraction times between the two simulated models the Wilcoxon signed-rank test was used. The values in the text for coherence results are reported as mean [95% confidence intervals]. All the other values are reported as median (interquartile interval) both in the text and in the figures.

All individual data of motor unit discharge times recorded at 90° and 130° are available at https://doi.org/10.6084/m9.figshare.24631191.

## Results

### Dorsiflexion torque output

To assess how changes in TA muscle length affected isometric dorsiflexion torque production, we compared the peak MVC and torque output steadiness between ankle joint angles (90° and 130°). Peak MVC values significantly decreased from 33.1 (28.9–35.9) Nm at 90° to 30.2 (25.6–33.2) Nm at 130° ankle joint angle (Wilcoxon signed-rank test; V = 86; *P* = 0.035). **Figure 2A** shows the variation in dorsiflexion torque output at the selected ankle joint angles (90° in blue and 130° in red) during the 10% MVC for a representative participant. The results observed for this representative participant were confirmed in the group data, showing no significant differences in torque output steadiness between 90° and 130° ankle joint angles (Wilcoxon signed-rank test; V = 46; *P* = 0.715). The coefficient of variation of torque output values were 2.1% (1.7–3.3%) and 2.4% (1.8–2.7%) for 90° and 130° ankle joint angles, respectively.

**Figure 2:**
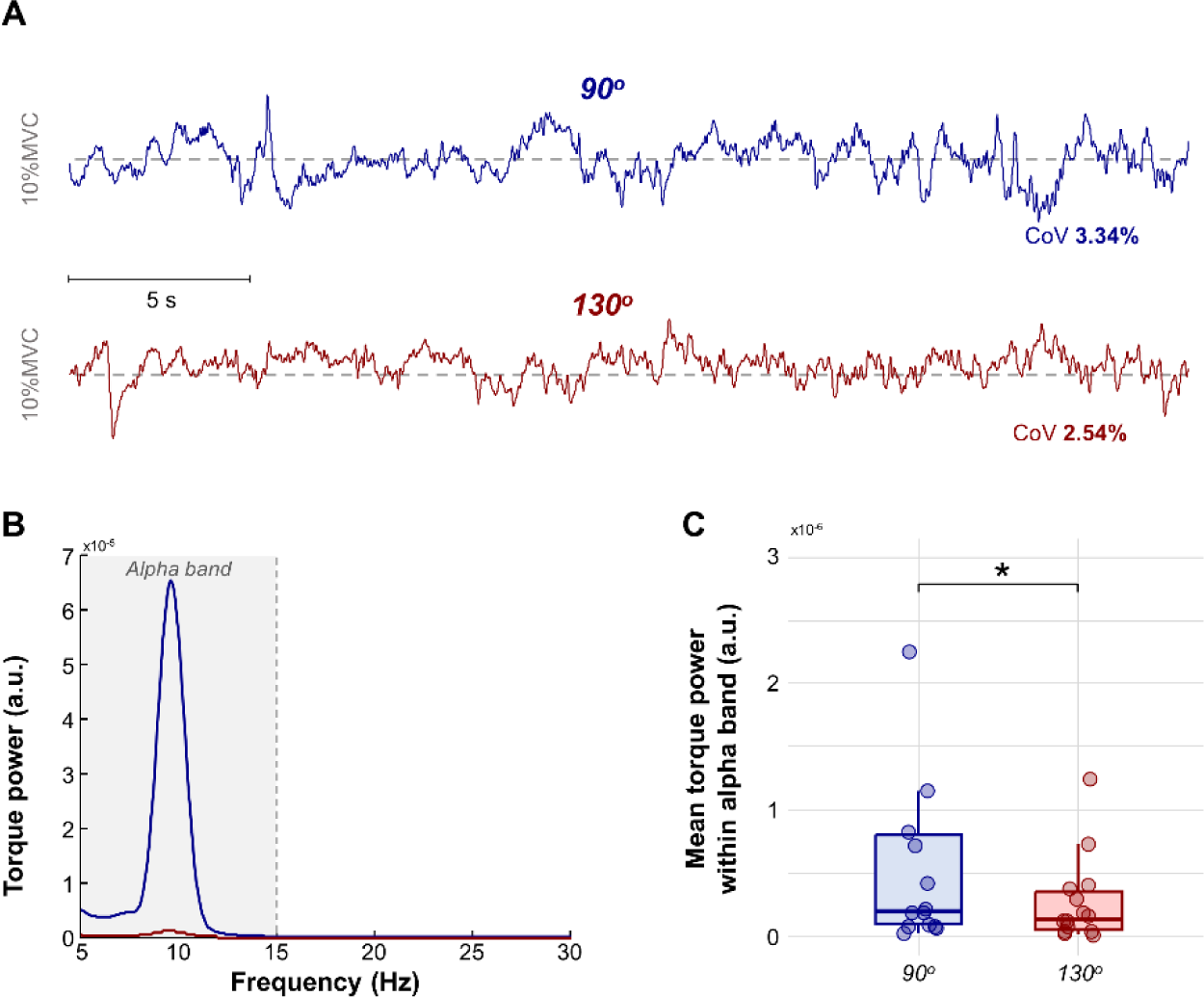
Torque output results. A, Torque fluctuations for a representative participant, the blue and red lines indicate the ankle joint angle at 90° and 130°, respectively. Note that the coefficient of variation (CoV) is similar between the torque signals. B, Power spectrum of torque signals depicted in A. The grey area shows the alpha frequency bandwidth (5-15 Hz). C, Group results of mean torque power within the alpha band. Circles identify individual participants. Horizontal traces, boxes, and whiskers denote median value, interquartile interval, and distribution range. *P<0.05.

We calculated the mean torque output power within the 5-15 Hz range to examine changes in torque output oscillations within the alpha band between the ankle joint angles. **Figure 2B** depicts the power spectrum of the torque output obtained for the same participant of **Figure 2A**. The power within 5-15 Hz is markedly higher when the ankle joint angle was at 90° than 130°. When considering the group results, mean torque output power values significantly decreased between 90° and 110° (**Figure 2C**; Wilcoxon signed-rank test; V = 93; *P* = 0.009).

### Level of common synaptic input

To evaluate changes in common synaptic input between conditions HDsEMG signals were decomposed into their constituent motor unit spike trains. Subsequently, the average of z-coherence estimates was then quantified in the delta (0-5 Hz), alpha (5-15 Hz) and beta (15-35 Hz) bands, separately for each ankle joint angle. A total of 225 and 215 motor units were included for the 90° and 130° ankle joint angles, respectively. All details regarding the number of motor units included for each participant are provided in **Table 1**.

**Table 1:** Number of motor units included for each participant.

| Participants | Ankle Joint Angle |  |
| --- | --- | --- |
|  | 90° | 130° |
| #1 | 9 | 12 |
| #2 | 6 | 4 |
| #3 | 13 | 3 |
| #4 | 18 | 16 |
| #5 | 19 | 3 |
| #6 | 17 | 18 |
| #7 | 18 | 19 |
| #8 | 22 | 19 |
| #9 | 31 | 32 |
| #10 | 23 | 25 |
| #11 | 35 | 29 |
| #12 | 18 | 12 |
| #13 | 17 | 14 |
| #14 | 11 | 15 |
| **Total** | **257** | **221** |
| **Median** | **18** | **16** |
| **Interquartile interval** | **14–21** | **12–19** |

Two participants (#3 and #5; **Table 1**) were excluded from the coherence analysis as only three motor units were decomposed at 130° (see *Methods*). **Figure 3** shows the z-coherence estimates for a representative participant. As the ankle joint angle increased (increasing the TA muscle length) there was a concomitant reduction in the coherence in the 5-15 Hz bandwidth (alpha band; see highlighted yellow area in **Figure 3**).

**Figure 3:**
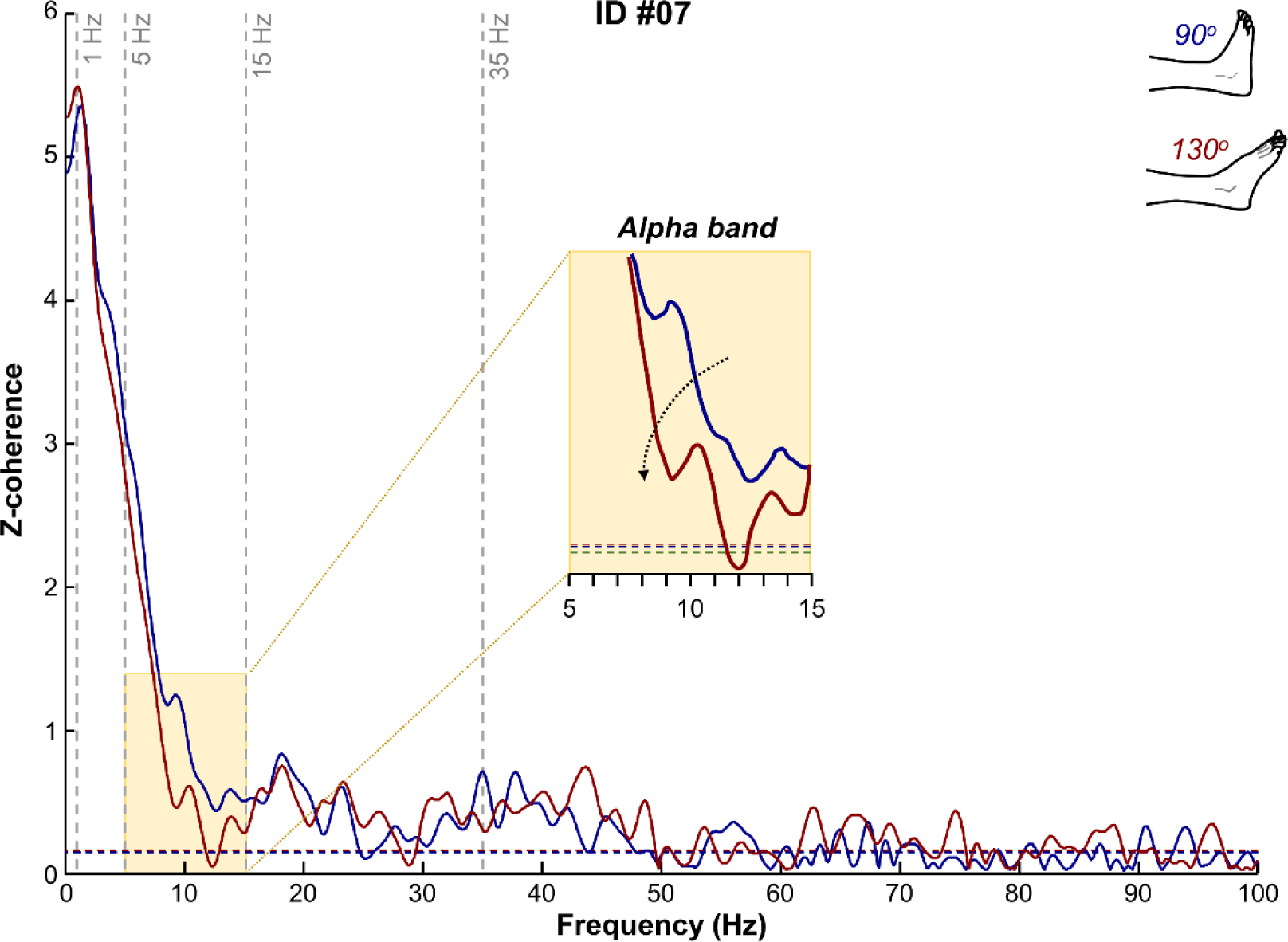
Representative result of motor units’ z-coherence. Motor units’ z-coherence profiles obtained for a representative participant (ID #07), where the blue and red lines indicate the ankle joint angle at 90° and 130°, respectively. The horizontal dashed line indicates the confidence level. Vertical dashed lines highlight the three frequency bandwidths analyzed: delta (1-5 Hz), alpha (5-15 Hz), and beta (15-35 Hz) bands. The yellow box depicts a zoom in of the alpha band.

Similar results were observed for the group data. **Figure 4A** depicts the pooled z-coherence considering all participants. Greater values of alpha z-coherence were obtained for the condition with the ankle joint angle at 90° compared to 130°. There was a significant effect of condition on the coherence values in the alpha band (**Figure 4C**; LMM; *F* = 24.64; *P* < 0.001), which was not observed in either the delta (**Figure 4B**; LMM; *F* = 0.69; *P* = 0.422) or beta bands (**Figure 4D**; LMM; *F* = 2.89; *P* = 0.117). Specifically, z-coherence values in the alpha band significantly reduced from 1.20 [0.95, 1.45] to 0.99 [0.73, 1.24] between 90° and 130° ankle joint angles.

**Figure 4:**
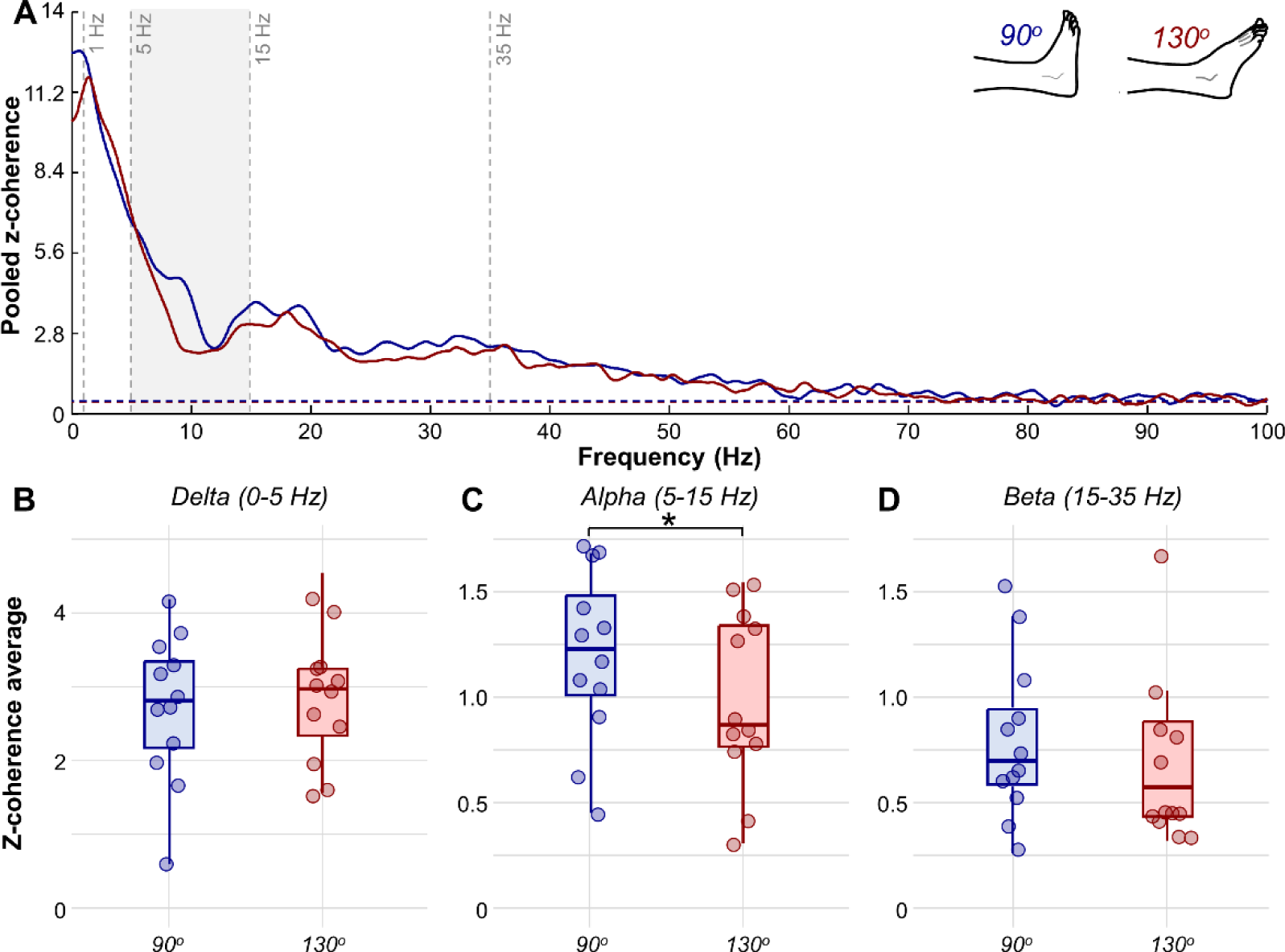
Z-coherence results. A, Pooled z-coherence profiles considering all participants (blue for 90° and red for 130°). The horizontal dashed line indicates the confidence level. Vertical dashed lines highlight the three frequency bandwidths analyzed: delta (1-5 Hz), alpha (5-15 Hz), and beta (15-35 Hz) bands. Grey area denotes statistical differences in the z-coherence between ankle joint angles. B-D, Group results of the average z-coherence for the delta (B), alpha (C), and beta (D) bands. Circles identify individual participants. Horizontal traces, boxes, and whiskers denote median value, interquartile interval, and distribution range. *P<0.05.

### TA twitch torque

To assess how changes in muscle length affected the evoked twitches from the TA, we conducted a second set of experiments in a subgroup of five participants where electrical stimuli were applied to the common peroneal nerve. This was repeated for the ankle positioned at 70° (shortened muscle length) and 130° (lengthened muscle length) to compare the extremities of the full range of dorsi-plantar-flexion motion. The individual results for the five participants are displayed in **Figure 5**. It is possible to qualitatively observe in the top panel of **Figure 5A** that the duration of TA evoked twitches increased for all participants when the ankle joint position was changed from shortened (yellow line) to lengthened (red line). *t_peak_* values were greater when the muscle was lengthened than at a shortened length for all participants, with increases ranging from 8.3 to 14.1 ms (see orange traces in the bottom panel of **Figure 5A**). *t_50%peak_* values also increased for all participants when comparing the lengthened to shortened conditions, with increases ranging from 29.2 to 54.2 ms (see green traces in the bottom panel of **Figure 5A**). Moreover, changing the ankle joint angle from 70° to 130° (i.e., increasing the TA muscle length) induced alterations in the low-pass filter characteristics of the TA. As can be observed in **Figure 5B**, the cut-off frequency *f_co_* (frequency in which the gain was attenuated by -3dB) was lower when the muscle was lengthened (red line) than shortened (yellow line) for all participants.

**Figure 5:**
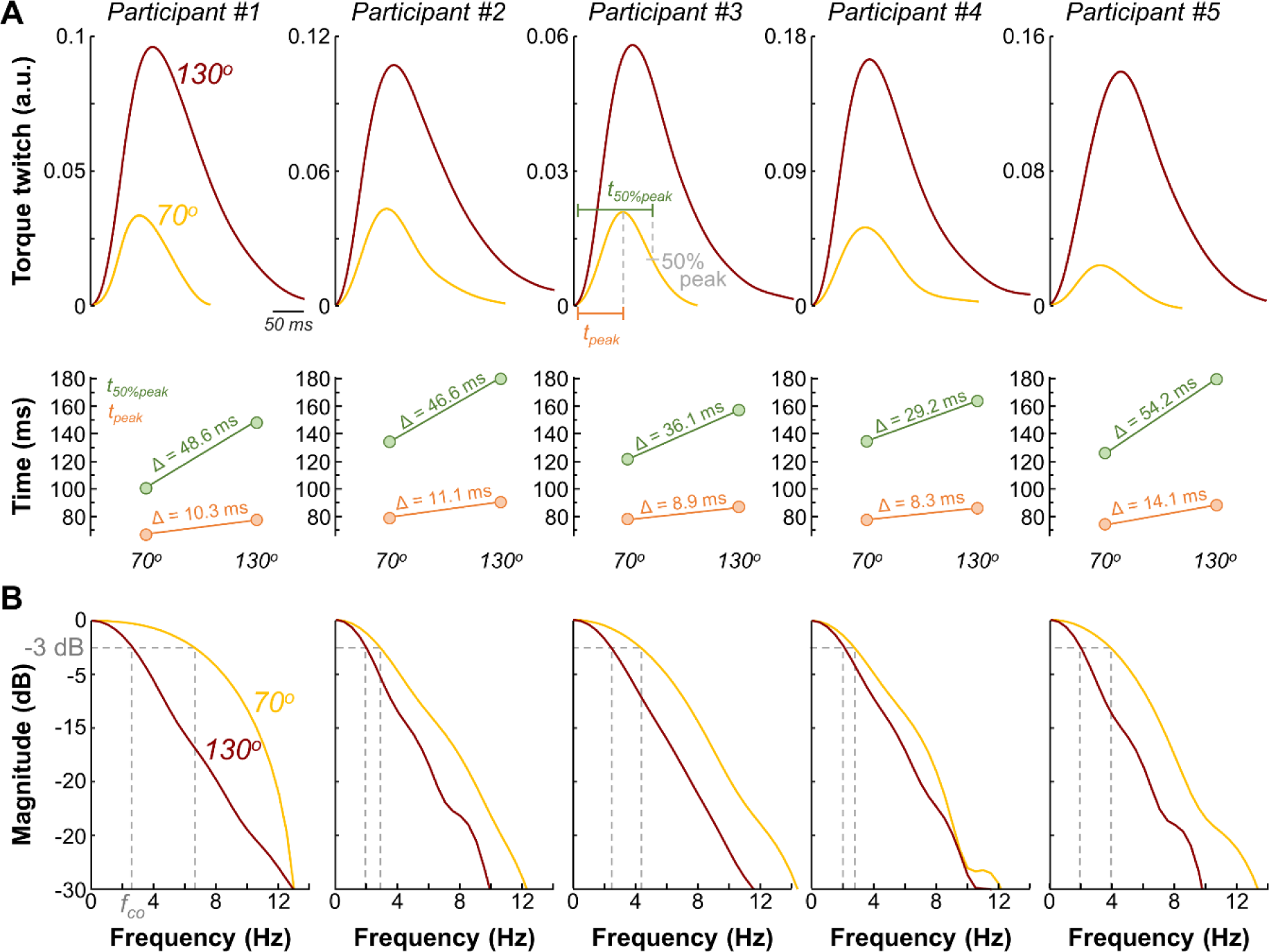
Twitch torque results. A, Individual results of twitch torque signals evoked from the tibialis anterior muscle when electrical stimuli was applied to the common peroneal nerve. Yellow and red lines indicate the ankle positioned at 70° (shortened muscle length) and 130° (lengthened muscle length), respectively. The bottom part of panel (A) shows the time to peak (t_peak_) and time to half-peak (t_50%peak_) changes when the ankle position was changed from shortened to lengthened. B, Magnitude of the frequency-response of the twitch torque obtained for each ankle joint angle (yellow for 70° and red for 130°). The cut-off frequency f_co_ in which the gain was attenuated by -3dB is indicated with the dashed grey lines.

### Simulations results

To explore how changes in the contractile properties of motor units affect the transmission of alpha band oscillations (physiological tremor) into muscle force, we simulated two models (model 1 and model 2) of ensembles of motor neurons receiving excitatory drive and generating isometric muscle force output. To simulate muscle lengthening conditions, we increased the twitch contraction times of individual motor units in model 2. We then calculated the z-coherence of motor units following the methodology described in the experimental data. **Figure 6A** illustrates the relationship between motor unit contraction time and motor unit twitch force output for the two generated models. Visually, we observe that model 2 (orange) exhibits a rightward shift towards greater twitch contraction times compared to model 1 (black). This difference, similar to that observed in the evoked twitch results, was confirmed statistically (**Figure 6B**; Wilcoxon signed-rank test; V = 0; *P* < 0.001). **Figure 6C** shows the pooled z-coherence for the ten realizations of each model and demonstrates a decrease in the z-coherence within the alpha band in model 2. Additionally, significantly lower z-coherence values were observed for model 2 compared to model 1 (**Figure 6D**; LMM; F = 16.84; *P* < 0.003). We also observed greater z-coherence values in the delta band (LMM; F = 95.71; *P* < 0.001), but no statistically significant differences were found in the beta band (LMM; F = 2.31; *P* = 0.163).

**Figure 6:**
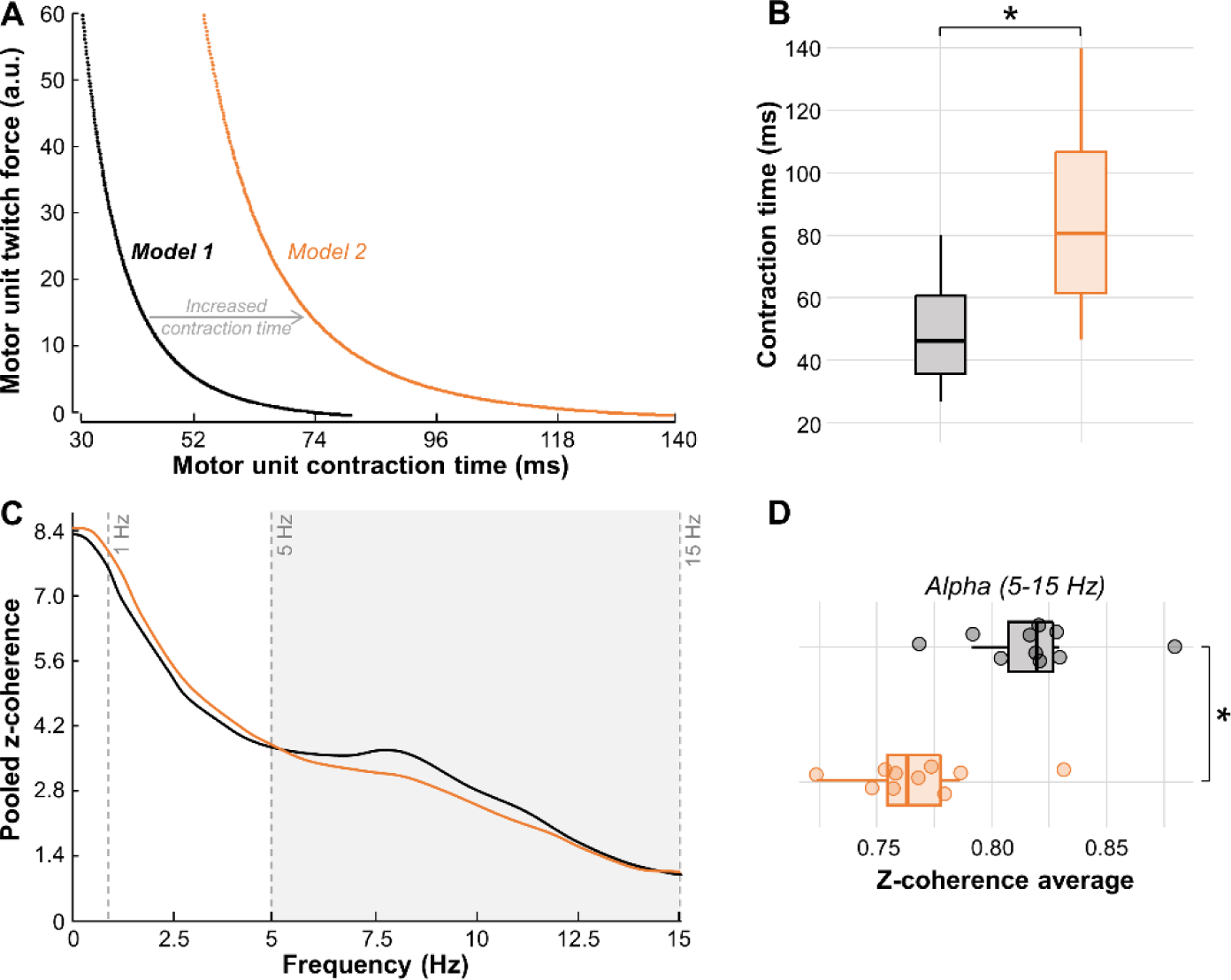
Simulation results. A, Relationship between motor unit contraction time and motor unit twitch force output for the two generated models of ensembles of motor neurons. Note that the increase in twitch contraction times of individual motor units in model 2 (orange) compared with model 1 (black). B, Comparison between motor unit contraction time of the two models. C, Pooled z-coherence profiles for the ten realizations of each model (black for model 1 and orange for model 2). D, Results of z-coherence values in the alpha band for the ten realizations of each model. Circles identify individual participants. Horizontal traces, boxes, and whiskers denote median value, interquartile interval, and distribution range. *P<0.05.

## Discussion

The present study investigates acute adjustments in neuromuscular control strategies based on changes in joint angle and consequently musculotendinous length. Our main findings revealed that changing the TA muscle length from shorter to longer induced reductions in the magnitude of alpha band oscillations in the common synaptic inputs. Additionally, we explored the relationship between these changes in physiological tremor and the low-pass filtering characteristics of the muscle. Our experimental results, supported by a simplified computational simulation, suggested that the increase of motor unit twitch duration resulting from increased muscle length directly influences the amplitude of the alpha band (physiological tremor) oscillations which may be through a proprioceptive (Ia) afferent loop.

### Modulation of alpha band oscillations in motor neuron inputs with changes in muscle length

During sustained isometric contractions, shared synaptic projections to the motor neuron pool generate significant coherence between motor unit spike trains across delta, alpha, beta, and even higher frequency bands. For instance, Castronovo et al. (2015a) have shown significant coherence up to ∼80 Hz during sustained isometric contractions, and these results have been consistently corroborated in subsequent studies using motor unit recordings (Muceli et al., 2022) and EMG measurements (Kerkman et al., 2018). However, due to the muscle’s contractile properties, which act as a low-pass filter on neural drive at ∼12 Hz (Bawa and Stein, 1976, Baldissera et al., 1998), only the coherent oscillations in the delta and alpha bands are transmitted to the force signal. Therefore, these oscillations are effectively responsible for force variability (Hug et al., 2023). Considering that delta band oscillations represent the precise command for optimal force generation (De Luca et al., 1982, Negro et al., 2009, Farina et al., 2014), alpha band oscillations can be viewed as involuntary common noise inputs that limit the accuracy of the force output (i.e., force steadiness). Consequently, modulations in alpha band coherence offer an interesting mechanism to modulate neural connectivity and enhance the precision of the force output. These modulations in alpha band have primarily been attributed to the resonant behaviour of Ia afferent pathways (Halliday and Redfearn, 1956, Lippold, 1971, Lippold, 1970, Cresswell and Löscher, 2000). Moreover, previous observations in macaque monkeys have suggested that specific spinal interneurons may cancel the alpha band frequency range from cortex to motor neuron inputs, thereby reducing tremor and improving movement precision (Williams et al., 2010, Koželj and Baker, 2014). Therefore, various factors, such as the visual demand of the task (Laine et al., 2014) or pain (Yavuz et al., 2015), can modulated the tremor frequency range in the common synaptic input to motor neurons, directly influencing optimal force output control.

Consistent with these studies, our results demonstrated that manipulating muscle length provides another approach to modulate physiological tremor in the shared synaptic inputs to the alpha motor neuron pool. Specifically, by increasing the TA muscle length, we observed significant reductions in the coherence between motor unit spike trains within the alpha band, while not significantly affecting the delta or beta bands (**Figure 4B-D**). These findings suggest that changing the muscle length from shorter to longer induces a reduction in the common synaptic noise input into motor neurons. Importantly, these reductions were reflected in the oscillations of muscle force output within the alpha band (**Figure 2C**), revealing that increasing muscle length offers a means to decrease physiological tremor frequency in force output and enhance motor control precision.

### Mechanisms involved in reducing tremor oscillations in the lengthened muscle

It has been reported that force output fluctuations due to changes in muscle length may stem from variations in the viscoelastic properties of the joint such as musculoskeletal stiffness and joint laxity or spindle Ia afferent gains and modulation in the motor unit discharge rate (Rack and Westbury, 1969, Clamann and Schelhorn, 1988, Powers and Binder, 1991, Jalaleddini et al., 2017, Inglis and Gabriel, 2021). Previous research has reported that higher stimulation frequencies are needed to achieve similar force outputs at shorter muscle lengths (Mela et al., 2001), implying the need for adjustments in motor unit discharge rates (Marsh et al., 1981, Rack and Westbury, 1969). However, during voluntary contractions, specifically for the TA, motor unit discharge rates remain stable across different muscle lengths (Bigland-Ritchie et al., 1992, Cudicio et al., 2022). This stability in the motor unit discharge rate suggests that other mechanisms, aside from the modulation of the motor unit discharge rate, might contribute to the alterations in force tremor as a result of changes in muscle length (Bigland-Ritchie et al., 1992). Considering the interplay between the Ia afferent loop and physiological tremor modulations (Lippold, 1970, Hagbarth and Young, 1979, Cresswell and Löscher, 2000, Christakos et al., 2006, Laine et al., 2016), Jalaleddini et al. (2017) employed a closed-loop model to investigate whether adjustments in Ia afferent pathway gain could explain changes in force tremor at various lengths of the plantar flexors. Jalaleddini et al. (2017) reported that increased tremor with muscle shortening could be attributed to adjustments of Ia afferent gains, particularly through gamma static fusimotor drive. To further explore these findings our study investigated another potential mechanism that could explain the reductions in tremor, both in the shared synaptic input to motor neurons and in force output, with muscle lengthening, that being the low pass filtering effect of the muscle. Consistent with previous findings (Mannard and Stein, 1973, Bawa and Stein, 1976, Bigland-Ritchie et al., 1992), we demonstrated that the twitch time to peak torque and the time to half of the peak torque were greater in the lengthened compared to shortened condition (**Figure 5A**). These results align with expected outcomes, as the decreased joint angle results in increased pre-contraction muscle tendon unit tension in the flexed (70°) condition which facilitates a faster achievement of peak twitch torque compared to the extended (130°) ankle joint position. Specifically, the ability to further shorten may be impeded by the already hyper-flexed ankle joint angle (70°), resulting in a lower twitch torque output achieved in a shorter period. The increased twitch duration with muscle lengthening enhances the low-pass filter characteristics of the TA (i.e., reduction of *f_co_* with muscle lengthening, **Figure 5B**), leading to alterations in the filtering of the neural drive to the muscle and, subsequently, in the Ia afferent feedback loop. Therefore, our results suggest that, in addition to fusimotor control (Jalaleddini et al., 2017), changes in the peripheral contractile properties of motor units due to alterations in muscle length have a significant impact on the transmission of the shared synaptic noise into muscle force output. These experimental findings were further supported by the simulation results, demonstrating that similar excitatory drives arriving at motor neuron ensembles with different twitch durations resulted in distinct alpha band coherence between motor unit spike trains (**Figure 6**).

Another potential mechanism that could explain our findings relates to the lengthening of the antagonist triceps surae muscle when the TA is in a shortened position. This lengthening of the antagonist muscle may lead to a reciprocal inhibition of the dorsiflexors during contractions as both a biomechanical and neural protection mechanism to avoid injury. While not specifically addressed in this study, it is well known that reciprocal inhibition can also influence the common synaptic input oscillations to motor neurons (Yavuz et al., 2018), potentially resulting in alterations in physiological tremor in the alpha band. Moreover, it is plausible that other neural pathways (both excitatory and inhibitory), such as inputs from Golgi tendon organs, joint receptors, and other somatic receptors, could contribute to this phenomenon.

### Effect of changes in muscle length on torque output

To evaluate the impact of varying TA muscle length on isometric dorsiflexion torque production, we compared peak MVC and torque steadiness across different ankle joint angles (90° and 130°). Our results revealed that when the ankle was secured at a joint angle of 90°, peak torque significantly increased compared to the 130° position (∼8.7%). The lower MVC values at 130° could be explained, at least in part, by sarcomere overlengthening beyond the optimal overlap (Bigland-Ritchie et al., 1992, Petrovic et al., 2022). Interestingly, in the current study there were no significant differences in torque steadiness between the 90° or 130° ankle joint positions. It may be the viscoelastic properties of the muscle tendon unit acting about the ankle joint that may explain why there were no changes in force steadiness at 130° compared to 90°. At a shortened tendon length (90°), there is a significant amount of slack in the viscoelastic properties of the muscle tendon unit, which must be compensated for. Force output from the contractile tissue must overcome the elasticity or slack of both the TA tendons (aponeurosis) and the actomyosin cross-bridge, known as the series and parallel elastic components (Kaufman et al., 1991), to generate smooth movement about a joint. In order to overcome this slack and generate tension down the tendons line of action it may require motor unit behavioural changes such as synchronization, doublet discharges (Yao et al., 2000, Enoka et al., 2003) or the recruitment of additional motor units (Farina and Negro, 2015). Unfortunately, this may come at the expense of force output stability (Inglis and Gabriel, 2021). Conversely, placing the joint at 130° significantly removes the slack in the system, optimizing the speed of tension transmission down the tendons line of action, resulting in a shorter electro-mechanical delay (Yavuz et al., 2010). However, this comes at the cost of stability in force output steadiness, as the lengthened position likely increases the number of active cross-bridges and, consequently, internal force (Ng et al., 1994, Wu et al., 2019), leading to a mechanical disadvantage for steady force production.

### Limitations

The positioning of participants during testing is a potential limitation of the study that may have influenced the results. In this study, subjects were seated with the test leg extended. Mitchell et al. (2008) reported that the range of motion of the ankle joint can be significantly inhibited based on the position of the pelvis, hip, and knee joints. Their study demonstrated that when the knee is flexed (90°) compared to extended (180°) in a seated position, the ankle joint’s passive range of motion changes from 47.3° to 16.4°, representing almost a 65% reduction in the range of motion. Changes in the hip and knee angles may either increase or decrease tension in the passive tissues. Therefore, straightening the leg while seated may passively increase tension in the nerves, muscles, and tendons from the lumbar spine to the foot, potentially influencing the range of motion of the ankle joint. However, the evident changes in peak torque among ankle joint angles suggest that we successfully investigated positions at both shortened and lengthened relative to an optimal length. Additionally, flexing the knee would naturally induce a shortening in the biarticular triceps surae muscle, which could be a potential confounding factor.

### Conclusions

Our study explored the relationship between muscle length alterations and neural changes affecting isometric dorsiflexion torque production. We observed that changing the TA muscle length from shorter to longer resulted in decreases in alpha band oscillations (physiological tremor) both in the shared synaptic input to motor neurons and in force output. Importantly, our findings suggest that alterations in the average motor unit twitch duration contribute to these observed effects, influencing the synaptic input to the motor neuron pools through the Ia afferent loop and, consequently, the neural drive to motor neurons. Therefore, the current study provides valuable insights into the interplay between muscle biomechanics and neural adjustments, while illuminating potential neuromechanical mechanisms underlying force production and control. These results may be utilized in either clinical populations or the elderly in the prescription of rehabilitation strategies to focus more on proprioceptive feedback during recovery (i.e., post-stroke, post-fall).

## Author contributions

CO, UY and FN contributed to study design; AC and MC acquired the data; HVC and FN analyzed the data; HVC, JGI and FN interpreted the data; HVC, JGI and FN drafted the manuscript; HVC, JGI, AC, MC, CO, UY and FN revised and edited the manuscript. All authors have approved the final version of the manuscript and agree to be accountable for all aspects of the work. All persons designated as authors qualify for authorship, and all those who qualify for authorship are listed.

## Acknowledgments

This study was funded by the European Research Council Consolidator Grant INcEPTION (contract no. 101045605).

